# Whitebait conservation and protected areas at non-tidal rivermouths: integrating biogeography and environmental controls on īnanga (*Galaxias maculatus*) spawning grounds

**DOI:** 10.1101/2020.11.06.372243

**Authors:** Shane Orchard, David R. Schiel

## Abstract

*Galaxias maculatus* is a declining amphidromous fish that supports culturally-important whitebait fisheries in New Zealand and elsewhere in the Pacific. As a largely annual species, the seasonal productivity of spawning grounds has a strong influence on the availability of recruits. Spawning ground protection is urgently required to reverse historical degradation and improve prospects for the maintenance of sustainable fisheries. Although spawning habitat has been well characterised in tidal rivers where it is structured by water level changes on spring high tides, there has been no previous study of spawning in non-tidal rivermouths. We assessed seven non-tidal rivers over four months using a census survey approach to quantify spawning activity, identify environmental cues, and characterise fundamental aspects of the biogeography of spawning grounds. We report conclusive results that include a) identification of compact spawning reaches near the rivermouths, b) triggering of spawning events by periods of elevated water levels that were often of very short duration, suggesting that potential lunar cues were less important and that rapid fish movements had likely occurred within the catchment prior to spawning events, and c) consistent vertical structuring of spawning grounds above typical low-flow levels with associated horizontal translation away from the river channel, leading to increased exposure to anthropogenic stressors and associated management implications for protecting the areas concerned. These consistent patterns provide a sound basis for advancing the management of non-tidal rivermouths. Attention to flood management, vegetation control, and bankside recreational activities is required and may be assisted by quantifying spawning ground biogeography. The identification of rapid responses to environmental cues deserves further research to assess implications for floodplain connectivity management to support fish movements in emphemeral flowpaths, and as a potential source of bias in commonly-used fish survey methodologies.

## 1. Introduction

*Galaxias maculatus* is a small-bodied amphidromous fish that is widespread across the Pacific where it is known by various names including īnanga, common jollytail, puye and puyen (Barile et al. 2015; McDowall 1991; Zattara & Premoli 2005). Traditional ‘whitebait’ fisheries targeting juvenile fish have been established in several countries including New Zealand, Australia, Chile and Argentina, all of which have suffered historical declines that impact the viability of continued harvest (Campos 1973; Fulton 2000; Mardones et al. 2008; McDowall 1984; Vega et al. 2013). These fisheries are situated near the coastal interface and exploit the post-larval stage as young fish migrate into freshwater systems following an oceanic development period of ca. 6 months (McDowall & Eldon 1980). At this stage, juvenile *G. maculatus* are largely colourless and typically in the 40–55 mm size range, though undergo rapid developmental changes soon after entering freshwater systems (McDowall 1968; McDowall et al. 1994). New Zealand’s fishery has considerable recreational, cultural and economic value that contributes to the wellbeing of local communities nationwide and can be particularly important in small towns and rural areas (McDowall 1984). Although *G. maculatus* is the most abundant species in the whitebait catch in the majority of rivers, four other migratory galaxiids (*G. argenteus, G. fasciatus, G. brevipinnis* and *G. postvectis*) also contribute various proportions, and in some cases augmented by a sixth non-Galaxiidae species *Retropinna retropinna* (common smelt) (McDowall 1965; Yungnickel et al. 2020).

The migratory lifecycle of these fishes has important implications for the existence and sustainability of whitebait fisheries at a variety of scales (McDowall 2008). At the sub-global scale, the general pattern of amphidromy involves an oceanic development period in which larvae may be transported long distances and potentially colonise distant shores (McDowall 2002, 2007). This life history may be evolutionarily favourable in small island states otherwise depauperate in freshwater fish fauna by providing a mechanism for recruitment to new habitats and leading to the establishment of widely distributed populations (Berra et al. 1996; McDowall 2010a). However, at more local scales the replenishment of larval sources remains critical to the maintenance of existing populations. In many cases this relies on the continued health of adult fish populations for egg production coupled with the bio-regional partitioning of larval pools leading to recruitment pulses, or absences, as the case may be (McDowall 2010b). In addressing these dynamics, there is considerable debate about the relative importance of oceanic dispersion pathways, larval retention effects, and potential for natal homing, all of which may affect the biogeography of amphidromous larval pools at a variety of scales (Augspurger et al. 2017; McDowall 2010a; Waters et al. 2000).

Hickford & Schiel (2016) investigated the topic of natal homing in New Zealand and found no evidence of juvenile *G. maculatus* returning to the same rivers in which they hatched in a study of 12 catchments on the South Island’s east and west coasts. However, some recruits were identified returning to rivers close to their natal origin on the same coast, and 15 juveniles originating from the west coast were identified in east coast whitebait samples (Hickford & Schiel 2016). These results reflect the prevailing oceanic currents and are consistent with an important role for current-mediated transport in the structuring of regional larval pools (Chiswell & Rickard 2011). However, several other lines of evidence also point to the importance of local recruitment sources for the maintenance of local populations at certain times. At one extreme, the capacity for a residential and largely non-migratory lifecycle has been identified through studies on land-locked water bodies in Australia (Chapman et al. 2006; Pollard 1971) and South America (Barriga et al. 2007; Barriga et al. 2002; Carrea et al. 2013; Cussac et al. 2004). Inherent consequences include larval retainment such that all phases of the life cycle are completed within a significantly smaller spatial domain in comparison to a diadromous lifecycle (McDowall 2010c; Rojo et al. 2020). There is also evidence for a wide spectrum of life histories somewhere between these extremes, such as where larvae drift relatively slowly downstream and complete variable proportions of their growth and development in retentive freshwater bodies (Augspurger et al. 2017), or nearshore coastal environments such as river plumes (Shiao et al. 2015; Sorensen & Hobson 2005).

For *G. maculatus*, the potential effects of larval retention and bio-regional partitioning are heightened in comparison to other whitebait species because of its shorter, generally annual life cycle (McDowall 2010b). Given the high rate of turnover, stability in yearly egg production and recruitment trends can be critical for the maintenance of sustainable populations (McDowall et al. 1994). An additional piece of this puzzle is presented by variance in the biogeography of the spawning grounds themselves, with local egg production sources become disproportionately more important to the availability of post-larval recruits where partitioning occurs. Repeat declines are potentially disastrous due to the immediacy of cause-effect relationships between egg production, recruitment, and population size.

Although the biogeography of *G. maculatus* spawning grounds has been relatively well characterised in tidal rivers where they are strongly influenced by spring high tides (Benzie 1968; Orchard et al. 2018a; Orchard et al. 2018b; Taylor 2002), many non-tidal rivers also support fish populations. In New Zealand, these rivers are particularly common on the east coast of the South Island and south-eastern North Island where they are often associated with mixed sand-gravel beaches at the coast (Kirk 1980, 1991). Many such rivers form perched lagoons or *hāpua* under the influence of high wave energies and associated barrier formation that promotes the establishment of non-estuarine conditions at the rivermouth (Hart 2009; Kirk 1991). Other examples are found in regions with significant alluvial transport from glaciated and paraglacial areas (e.g., Iceland, Russia, Alaska and Canada), and also on fine-grain shorelines in Israel and South Africa (Carter et al. 1989; Cooper 1994; Green et al. 2013; Lichter & Klein 2011; McSweeney et al. 2017; Smith et al. 2014). To date, however, there has been no comprehensive assessment of the location, timing and environmental controls on *G. maculatus* spawning activity at non-tidal rivermouths.

The present study addresses this information gap in a region characterised by a high density of non-tidal rivermouths which facilitated both concurrent investigations at several study sites and the generation of a wider landscape view. Specific hypotheses included that spawning grounds were likely to be found in rivermouth lagoon environments due to similarities with tidal lagoons including their ability to support periodically inundated riparian vegetation, and that lunar cycles associated with spring high tide periods were unlikely to act as a trigger for spawning activity, as is the case in tidal rivers, due to the apparent lack of appreciable hydrodynamic effects. Instead, we considered that rain-induced water level variations would be more likely to trigger spawning events as has been reported in land-locked systems (Chapman et al. 2009; Pollard 1971) and tidal lakes (Richardson & Taylor 2002). Despite these expectations, the research was largely exploratory and involved the characterisation of previously unsurveyed rivers and their associated environments. In the remainder of the paper we report conclusive results that include strong spatiotemporal patterns likely to be transferable to other localities. These provide a sound basis for advancing the management of non-tidal rivermouths to support their role in whitebait conservation.

## 2. Methods

### 2.1 Study area

The Kaikōura coast is characterised by a predominance of non-tidal rivers as is the case elsewhere in the Canterbury region of New Zealand (see Suppl. Material Fig. S1). Seven non-tidal waterways of varying character were chosen within a ~35 km section of coastline between Oaro and Mangamaunu Bay (Fig. 1). These study catchments included two relatively large rivers (Oaro and Kahutara) and five smaller streams. Within this area there were no previous records of *G. maculatus* spawning grounds in the National Īnanga Spawning Database (NISD) or other known sources prior to this study. However, records in the New Zealand Freshwater Fish Database (NZFFD) include *G. maculatus* observations in five of the seven study catchments, and further populations were expected to be present in other waterways (see Suppl. Material Fig. S2). Whitebait fishing occurs in at least three of these rivers (Oaro, Kahutara and Lyell Creek).

**Fig. 1.**
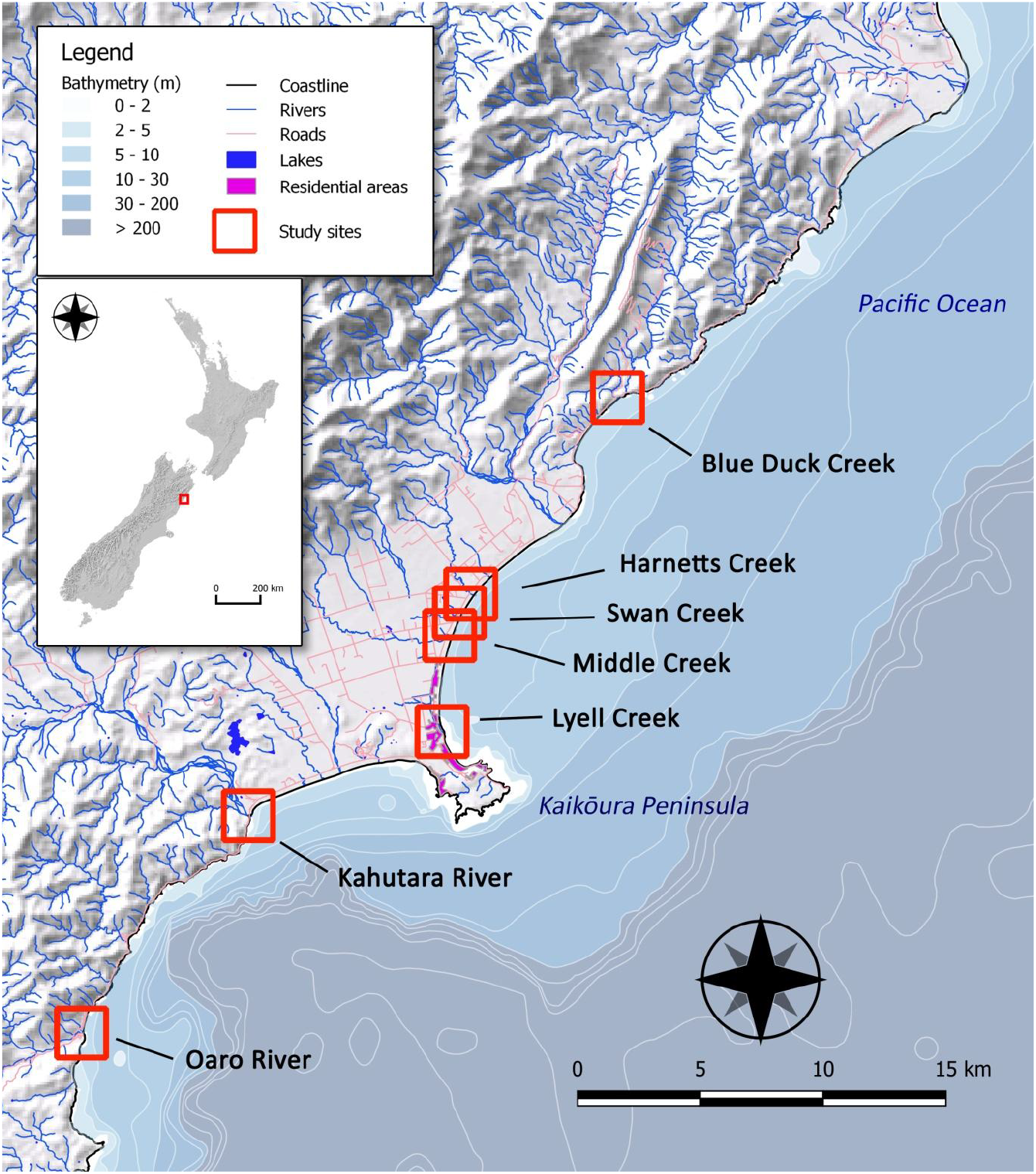
Location of study sites on the Kaikōura Coast, South Island, New Zealand.

The study catchments range from 440 ha to over 23, 000 ha in size with the smallest (Blue Duck) representing a 2^nd^ order stream originating in coastal hill-country, and the largest (Kahutara) being a 5^th^ order river originating in the Seaward Kaikōura Range (Table 1). Predominant catchment geologies partly reflect the inland extent of each catchment in relation to hill-country and coastal outwash plains, with the latter being extensive either side of Kaikōura Peninsula where they form mixed sand-gravel beaches at the coast (Fig. 1). As is characteristic of many waterways in the region, five of the study sites featured non-estuarine lagoons at the river mouth. At Blue Duck and Harnetts creeks no such lagoon was present during the study period, but they have been recorded in the past. All of the study sites, with the exception of Oaro, were affected by tectonic uplift of between 0.6 – 0.9 m during the 2016 Kaikōura earthquake (Clark et al. 2017; Schiel et al. 2019), which has generally increased the elevation of these lagoons and lower river reaches in relation to sea level. In the case of Lyell Creek, high spring tides would occasionally inundate the lower reaches pre-earthquake if combined with significant swell, but this no longer occurs since the uplift event (P. Adams, pers. comm.).

**Table 1.** Characteristics of study catchments as recorded in field observations and selected data from the New Zealand River Environment Classification (Snelder & Biggs 2002).

| Study catchment | Order† | Catchment area (ha) | Climate | Geology | Land cover | Salinity (ppt)‡ | Source of flow |
| --- | --- | --- | --- | --- | --- | --- | --- |
| Oaro River | 4 | 4725 | Cool wet | Hard sedimentary | Scrub | 0.01 | Hill country |
| Kahutara River | 5 | 23095 | Cool wet | Hard sedimentary | Scrub | 0.01 | Hill country |
| Lyell Creek | 3 | 1633 | Warm dry | Alluvium | Pastoral | 0.01 | Low elevation |
| Middle Creek | 4 | 2588 | Cool dry | Alluvium | Pastoral | 0.01 | Low elevation |
| Swan Creek | 3 | 1541 | Cool wet | Alluvium | Pastoral | 0.01 | Low elevation |
| Harnetts Creek | 3 | 1201 | Cool wet | Alluvium | Pastoral | 0.01 | Low elevation |
| Blue Duck Creek | 2 | 440 | Cool wet | Soft sedimentary | Scrub | 0.02 | Hill country |

† stream order of the lowest reach
‡ maximum bottom salinity recorded in the rivermouth lagoon during the study period

### 2.2 Spawning ground surveys

#### 2.2.1 Sampling design

In the first month of the study (February) intensive searches were conducted in all catchments beginning at the rivermouth and working upstream with the distribution of spawning sites (if any) being unknown. Although March and April were expected to be the months of peak spawning activity based on previous work in tidal waterways in the Canterbury region (Orchard et al. 2018a; Orchard et al. 2018b), significant spawning activity was detected in February that was apparently triggered by a rain event. This finding helped to establish the general location of spawning in the study catchments and define areas for repeat surveys in the following months. This resulted in the delineation of survey reach lengths of between 400 and 700 m from the rivermouth, with the upstream limit marked by a prominent riffle or general increase of gradient in the riverbed. None of the spawning sites detected in following months were located near the upstream limit of these survey areas, improving confidence that they were sufficient to detect the spawning activity that occurred. Data from permanent stage-height loggers operated by the Canterbury Regional Council were available for the Lyell and Middle Creek catchments (Environment Canterbury 2020). In both cases, these loggers were located several hundred metres upstream of the identified spawning reach but nonetheless provided a useful indication of water level fluctuations in relation to the timing of spawning events. Following discovery of the February spawning sites, additional water level loggers (Odyssey ODYWL, Dataflow Systems Ltd) were installed within the observed or potential/estimated spawning reach in the five remaining catchments to provide further information on water level changes in relation to subsequent spawning events. For all spawning sites detected, real-time kinematic (RTK) GPS surveys were conducted using a Trimble R8 GNSS receiver to measure the upper and lower elevation of the egg band. Geodetic benchmarks were included within all RTK-GPS surveys to achieve an estimated vertical accuracy of 3.5 mm + 0.4 ppm RMS. Heights were referenced to the New Zealand Vertical Datum (NZVD) (Land Information New Zealand 2016).

#### 2.2.2 Detection and measurement of spawning sites

##### Field surveys

Spawning surveys were completed over four consecutive months (February - May). Following the standard approach in tidal waterways in which spawning events have an apparent relationship with the lunar cycle (Orchard & Hickford 2018; Taylor 2002), each round of surveys commenced a few days after the full moon of the month, which was associated with the largest monthly spring tides within the study period (Table 2). The field survey technique involved systematic searches for eggs laid in riparian vegetation. For every 5 m length of river bank three searches were made at random locations. Each search involved inspection of the stems and root mats along a transect line oriented perpendicular to the river bank. Typically a 0.5 m wide swathe of vegetation was inspected on each transect, with these being of variable length depending on the bank slope and vegetation extent. In this case, the transect lengths were adjusted to include all vegetation within areas inundated by the February flood event as judged by the position of strand lines and silt deposition. As this was the highest water level recorded during the study period, the spatial extent established in February was applied to all subsequent surveys.

**Table 2.**
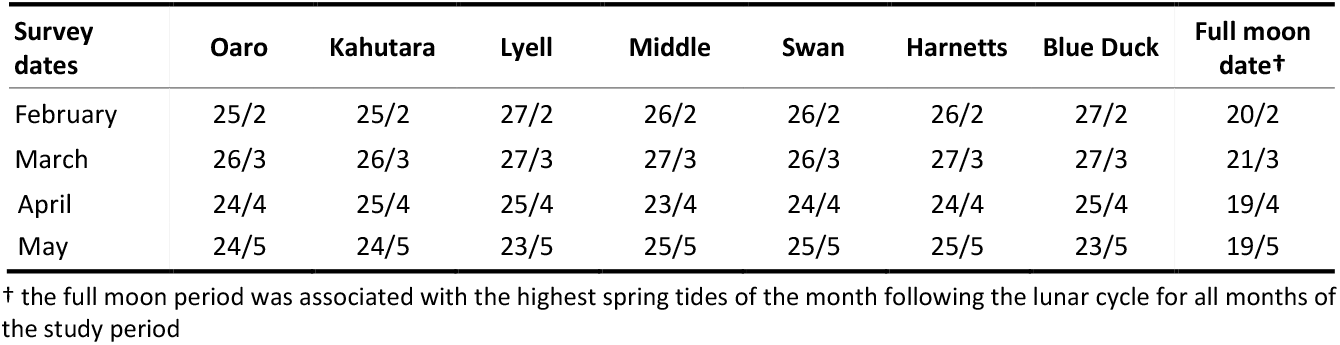
Survey dates and lunar cycle (day/month).

| Survey dates | Oaro | Kahutara | Lyell | Middle | Swan | Harnetts | Blue Duck | Full moon date† |
| --- | --- | --- | --- | --- | --- | --- | --- | --- |
| February | 25/2 | 25/2 | 27/2 | 26/2 | 26/2 | 26/2 | 27/2 | 20/2 |
| March | 26/3 | 26/3 | 27/3 | 27/3 | 26/3 | 27/3 | 27/3 | 21/3 |
| April | 24/4 | 25/4 | 25/4 | 23/4 | 24/4 | 24/4 | 25/4 | 19/4 |
| May | 24/5 | 24/5 | 23/5 | 25/5 | 25/5 | 25/5 | 23/5 | 19/5 |

† the full moon period was associated with the highest spring tides of the month following the lunar cycle for all months of the study period

##### Area of occupancy (AOO)

All egg detections were associated with a given location that was identified as a spawning site (Orchard & Hickford 2018). Individual sites were defined as continuous or semi-continuous patches of eggs with dimensions defined by the pattern of occupancy. In each case, the upstream and downstream extents of the patch were established and spawning sites coordinates were recorded using a hand-held GPS in the field and corrected in QGIS v3.12 (QGIS Development Team 2020) with the assistance of site photographs and landmarks. For each spawning site, the length along the riverbank was measured along with the width of the egg band at the position of each of the original search transect, and additional transects where needed to provide a minimum of three measurements at all sites. Zero counts were recorded when they occurred within a spawning site as is common where the egg distribution is not a continuous band. AOO was calculated as length x mean width.

##### Spawning site productivity

Egg production was assessed by direct eggs counts using a sub-sampling method (Orchard & Hickford 2016, 2018). At each transect, as above, a 10 x 10 cm quadrat was placed in the centre of the egg band and all eggs within the quadrat counted. Egg numbers in quadrats with high egg densities (>200 / quadrat), were estimated by further sub-sampling using five randomly located 2 x 2 cm quadrats and the average egg density of these sub-units used to calculate an egg density for the larger 10 x 10 cm quadrat. The mean egg density was calculated from all 10 x 10 cm quadrats sampled within the site, inclusive of zero counts. The productivity of each site was calculated as mean egg density x AOO.

### 2.3 Statistical analyses

Differences in AOO and egg production trends were tested using two-way ANOVAs with fixed factors of catchment and month, followed by post hoc Tukey HSD tests for significant main effects. Model assumptions were checked via Shapiro-Wilk and Levene tests using the *car* package in R (Fox & Weisberg 2011). Variation in AOO and egg production was also examined using robust one-way ANOVAs (Field & Wilcox 2017) to assess the effect of catchment and month separately using the *robust* package in R (Wang et al. 2020). These analyses considered catchment effects in relation to the pooled data from all months, and also for the two main spawning events treated separately (February and April). Similar tests were applied for the dependent variables of distance upstream, and vertical elevation based on the centrepoint of each spawning site in the horizontal and vertical dimensions, respectively. A regression analysis was used to explore the relationship between egg production and AOO using pooled data from all sites. All analyses were completed in R version 3.5.3 (R Core Team 2019).

## 3. Results

### 3.1 Occurrence, size and productivity of spawning grounds

Spawning sites were detected in all seven catchments, although at different times over the study period (Fig. 2). Two natural spawning events were responsible for the majority of spawning activity, the first occurring in February at the beginning of the study period, and the second in April. A total of 34 individual spawning sites were measured in these two events (14 and 20 sites, respectively). A single site was detected in March in one of the study catchments (Lyell Creek). However, this spawning event was triggered by the artificial raising of water levels as part of an ecological engineering experiment conducted with the local council, and was thus atypical of the wider regional trend. No spawning was detected in any of the study catchments in May.

**Fig. 2.**
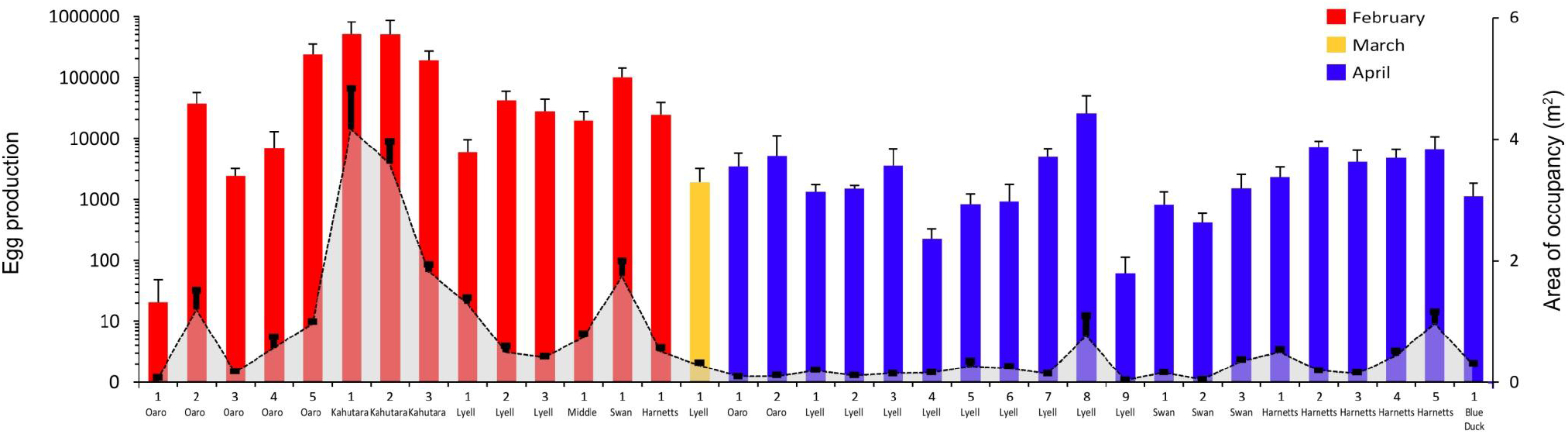
Summary of īnanga (*Galaxias maculatus*) spawning activity in seven non-tidal rivers on the Kaikōura Coast of New Zealand in 2019. Note log scale on Y-axis for egg production (columns) and linear scale for area of occupancy on right. Error bars are standard error of the mean derived from the sub-sampling approach used to calculate each metric for each spawning site. The X-axis shows individual spawning sites numbered by river for each of the three months in which spawning occurred. A fourth month (May) was also surveyed but no spawning was found.

Most of the spawning sites were relatively small though often dense patches of eggs. The total area of occupancy (AOO) recorded over all months (23.4 m^2^) was significantly different between rivers (*df* = 6, *F* = 13.71, *p* < 0.001), and similar results were obtained when these analyses were restricted to results from February and April (Table 3). The mean egg density across all sites and months was 3.8 eggs cm^-2^. A regression on the combined data showed a significant relationship between the AOO and egg production of individual spawning sites (*R*^2^ = 0.86, *F* = 189.1, *p* < 0.001). Decomposition to the individual months showed that the February relationship was stronger (*R*^2^ = 0.85, *F* = 69.38, *p* < 0.001) than in April (*R*^2^ = 0.34, *F* = 9.28, *p* = 0.007) reflecting the influence of the large and highly productive Kahutara River sites in February, but also the observation of relatively few sparse sites with low egg densities in relation to AOO.

**Table 3.** Summary statistics for īnanga (*Galaxias maculatus*) spawning grounds in seven non-tidal rivers on the Kaikōura Coast of New Zealand in 2019. AOO = area of occupancy. NZVD = New Zealand Vertical Datum 2016.

| Metrics |  | Oaro | Kahutara | Lyell | Middle | Swan | Harnetts | Blue Duck | All sites | df | Robust F | p-value | signif† |
| --- | --- | --- | --- | --- | --- | --- | --- | --- | --- | --- | --- | --- | --- |
| AOO (m²) | mean | 3.15 | 9.59 | 4.57 | 2.31 | 0.74 | 2.76 | 0.28 | 23.4 | 6 | 13.71 | 0.000162 | *** |
|  | SE | 0.60 | 1.15 | 0.75 | 0.29 | 0.06 | 0.40 | 0.04 | 3.3 |  |  |  |  |
| Egg Production | mean | 283304 | 1198965 | 116321 | 19092 | 101930 | 52634 | 1134 | 1773380 | 6 | 20.42 | 4.13E-06 | *** |
|  | SE | 181943 | 698446 | 86113 | 11150 | 73138 | 32977 | 715 | 745600 |  |  |  |  |
| Distance inland (m) | mean | 89 | 212 | 223 | 150 | 236 | 222 | 280 | 202 | 6 | 45.02 | 8.08E-12 | *** |
|  | SE | 6 | 22 | 19 | 0 | 28 | 16 | 0 | 13 |  |  |  |  |
| Elevation NZVD (m) | mean | 1.26 | 1.73 | 0.89 | 1.95 | 2.52 | 3.98 | 2.45 | 2.11 | 6 | 281.9 | 2.20E-16 | *** |
|  | SE | 0.04 | 0.05 | 0.03 | 0.05 | 0.06 | 0.11 | 0.03 | 0.05 |  |  |  |  |
| Number of sites |  | 7 | 3 | 13 | 1 | 4 | 6 | 1 | 35 |  |  |  |  |

| Metrics |  | Oaro | Kahutara | Lyell | Middle | Swan | Harnetts | Blue Duck | Contrasts | df | Robust F | p-value | signift |
| --- | --- | --- | --- | --- | --- | --- | --- | --- | --- | --- | --- | --- | --- |
| AOO (m<sup>2</sup>) | Feb | 3.0 | 9.6 | 2.2 | 1.8 | 0.7 | 0.5 | 0.0 | catchment | 6 | 15.34 | 5.13E-07 | *** |
|  | Apr | 0.2 | 0.0 | 2.1 | 0.6 | 0.0 | 2.3 | 0.3 | month | 1 | 4.55 | 2.98E-02 | * |
| Egg production | Feb | 274644 | 1198965 | 74850 | 19092 | 99167 | 24251 | 0 | catchment | 6 | 14.24 | 9.82E-07 | *** |
|  | Apr | 8660 | 0 | 39582 | 0 | 2763 | 28384 | 1134 | month | 1 | 8.89 | 0.00545 | ** |
| Distance inland (m) | Feb | 91 | 212 | 238 | 150 | 180 | 185 | na | catchment | 6 | 53.26 | 1.03E-13 | *** |
|  | Apr | 83 | na | 200 | na | 254 | 229 | 280 | month | 1 | 2.28 | 0.124 | ns |
| Elevation NZVD (m) | Feb | 1.29 | 1.73 | 0.96 | 1.95 | 2.53 | 3.52 | na | catchment | 6 | 273.3 | 2.20E-16 | *** |
|  | Apr | 1.20 | na | 0.86 | na | 2.52 | 4.08 | 2.45 | month | 1 | 0.18 | 0.667 | ns |
| Number of sites | Feb | 5 | 3 | 3 | 1 | 1 | 1 | 0 |  |  |  |  |  |
|  | Apr | 2 | 0 | 9 | 0 | 3 | 5 | 1 |  |  |  |  |  |

† significance codes: \*\*\* $p < 0.001$ , \*\* $p < 0.001$ , \* $p < 0.01$ , ns = not significant, $\alpha = 0.05$

The total egg production across catchments and months was ca. 1.8 million eggs (Table 3). Egg production differed significantly between catchments over the study period (*df* = 6, *F* = 20.42, *p* < 0.001) reflecting the influence of relatively high egg production at Kahutara in comparison to the other catchments, even though spawning was only recorded there in February (Table 3b). The between-month comparisons also showed significant differences for both AOO (*df* = 1, *F* = 4.55, *p* = 0.03), and egg production (*df* = 6, *F* = 8.89, *p* = 0.005) that is indicative of much greater spawning activity in February across the region as a whole. Two-way ANOVAs also showed significant differences between catchments (*df* = 6, *F* = 15.34, *p* < 0.001) and between months (*df* = 1, *F* = 7.80, *p* = 0.01), with no significant interaction between the two (*df* = 4, *F* = 1.42, *p* = 0.26).

### 3.2 Catchment position

All of the spawning sites were located surprisingly close to the coast, despite the presence of apparently suitable habitat for spawning further upstream (Fig. 3). This pattern held in all catchments with the mean upstream limit of spawning being 248 m from the rivermouth. The maximum distance upstream of any spawning site was 388 m in Lyell Creek, and the minimum distance ranged from 65 m to 275 m upstream of the rivermouth. This resulted in the identification of well-defined and relatively compact spawning reaches in all catchments by the conclusion of the study period (Fig. 3). These spawning reaches ranged from 5 m in length (associated with a single spawning site), to 251 m at Lyell Creek which also had the highest number of individual spawning sites (Table 2).

**Fig. 3.**
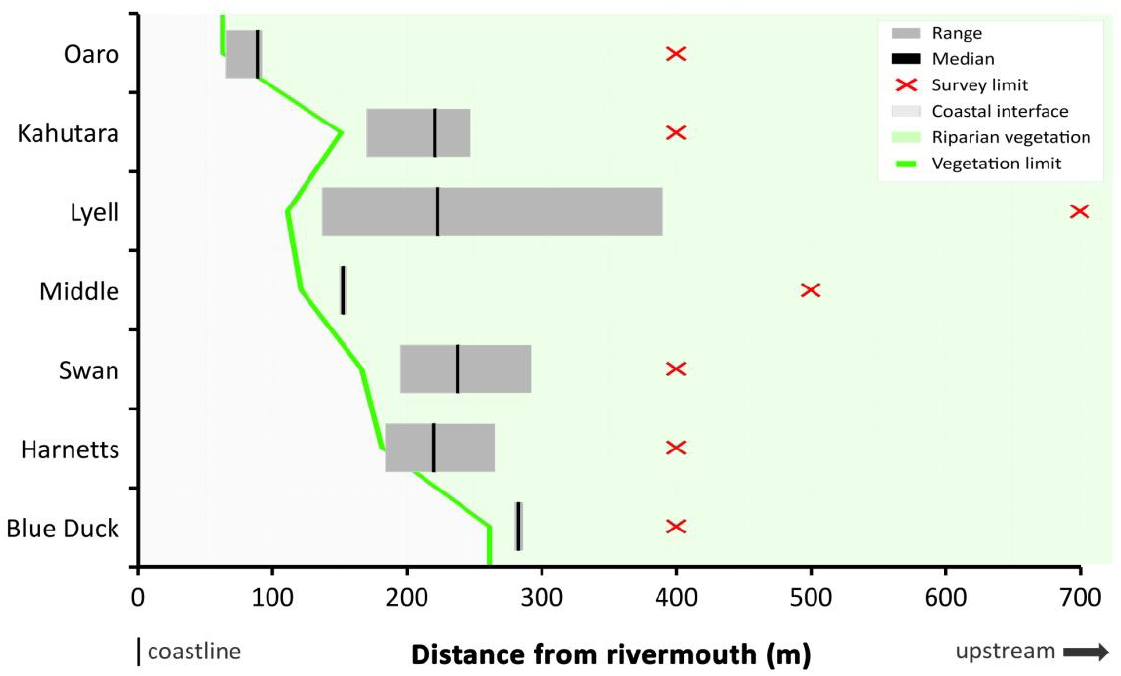
Horizontal position of īnanga spawning ground in seven non-tidal rivers on the Kaikōura Coast expressed as the range and median distance of the spawning reach upstream from the river mouth. The survey area extended from the coastline to the upstream limit indicated by red crosses. The green line indicates the downstream limit of riparian vegetation within the river channel as observed during the survey period. Seaward of this position all rivers are characterised by active mixed-sand gravel and boulder river mouth systems that are subject to regular reworking in storm and flood events.

The downstream limit of spawning also exhibited a close relationship with the downstream limit of riparian vegetation that was subject to at least periodic inundation during the study period (Fig. 3). This relationship was consistent between spawning events in the four study catchments where spawning was recorded more than once, as well as being common to all seven rivers despite marked differences in their morphology and size. The apparent high fidelity to a favoured spawning reach can be seen in the statistical comparisons which show a significant difference between catchments (*df* = 6, *F* = 45.0, *p* < 0.001) for the mean position of spawning sites expressed as distance from the rivermouth (Table 2a). This fidelity is partially obscured in the seasonal totals visualised in Fig. 3, but can clearly seen in the between-month comparisons for the two main spawning events (Table 2b). Differences in the distance inland were statistically significant between catchments (*df* = 6, *F* = 53.3, *p* < 0.001), but not between months (*df* = 1, *F* = 2.28, *p* = 0.124).

### 3.3 Vertical position

Significant differences were found between rivers in the vertical dimensions of spawning grounds. The highest elevation sites were in Harnetts Stream up to 4.5 m NZVD (New Zealand Vertical Datum 2016), and the lowest elevation sites (0.7 m NZVD) were in Lyell Creek (Fig. 4). In the other rivers spawning grounds fell within the range of 1.0 – 2.5 m NZVD, with the mean height across all individual spawning sites being 2.1 m ± 0.05(SE). For comparison, the Mean High Water Springs (MHWS) elevation on the Kaikōura coast was around 0.4 m NZVD within this period (LINZ 2018a, 2018b). One-way ANOVAs showed that the mean elevation of spawning sites was significantly different between sites across all months (*df* = 6, *F* = 273.3, *p* < 0.001), and similarly when only February and April were considered (Table 2a, b). However, mean elevation was not significantly different between February and April (*df* = 1, *F* = 0.18, *p* = 0.667). This reflects the occurrence of spawning sites in similar three-dimensional locations between months at Oaro, Lyell and Swan and Harnetts despite the addition of higher elevation sites at the latter in comparison to February (Fig. 4). These impressions are supported by two-way ANOVA results showing significant differences between catchments (*df* = 6, *F* = 458.2, *p* < 0.001), but not months (*df* = 1, *F* = 0.15, *p* = 0.669), and a significant interaction between the two (*df* = 3, *F* = 0.19, *p* = 0.001).

**Fig. 4.**
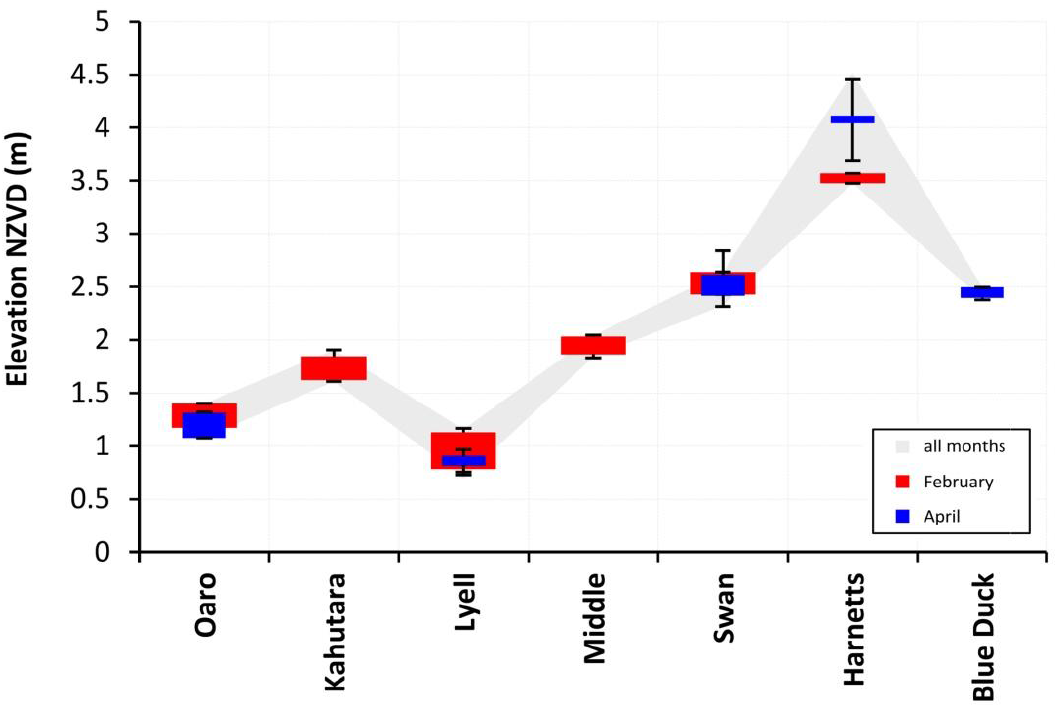
Vertical position of īnanga spawning ground in seven non-tidal rivers on the Kaikōura Coast. Coloured boxes represent the mean elevation of the egg band calculated from all sites recorded in the respective months. Error bars are one standard deviation of the mean position for top and bottom elevation of the egg band. The grey shading (all months) shows the maximum and minimum height of egg band in each river over all survey months. Note that the single spawning site recorded in March (in Lyell Creek) lies within the elevation band of other months.

### 3.d Role of environmental cues

There was a strong relationship between water level fluctuations and the timing of spawning events. The first of these occurred unexpectedly and involved an unseasonal heavy rain event in late February (Fig. 5a). Subsequent spawning surveys showed that this event triggered widespread spawning activity through the region that contributed 76% of the AOO and 95% of the egg production measured in the study period as a whole. Stage height data showed that the 23 February rain event caused a rapid spike in water heights of 70 cm in Lyell Creek, and 35 cm in Middle Creek (Fig. 5a), with this difference partly reflecting channel morphology and associated constriction effects at the logger positions as well as catchment-specific discharge rates. The apparent triggering effect of water level fluctuation had a considerable influence on the three-dimensional geography of the spawning grounds, with the lowest elevation eggs being located around 30 cm above typical low-flow levels in the spawning reach at all sites.

**Fig. 5.**
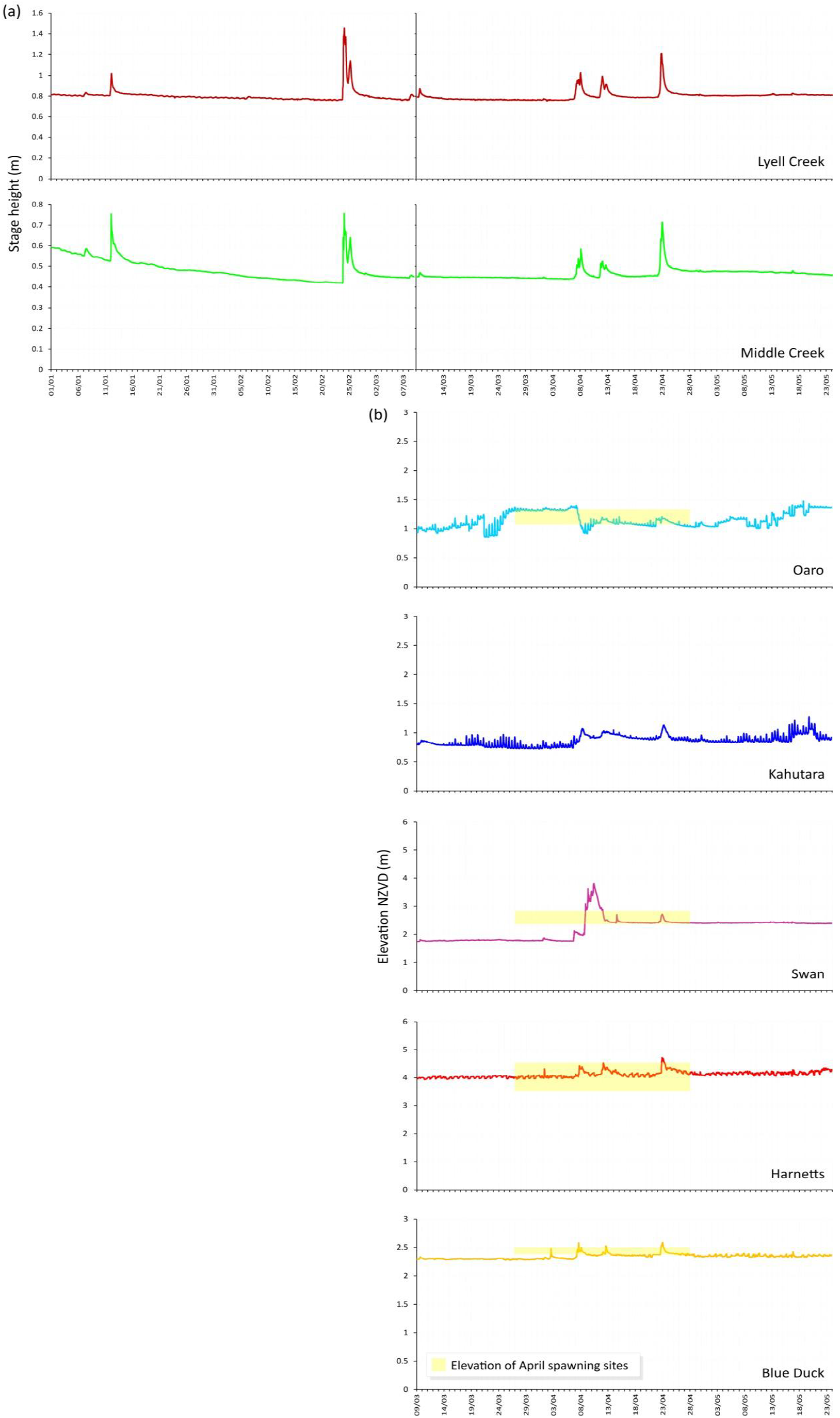
Hydrographs from the study catchments. (a) Stage height records from permanent loggers operated by the Canterbury Regional Council in Lyell Creek and Middle Creek for the summer of 2019. Each plot is split into two panels to facilitate comparison with the other catchments below. (b) Hydrographs from water level loggers deployed within the spawning reach of five catchments after the discovery *G. maculatus* spawning sites near the rivermouths in February. Note that these plots are referenced to New Zealand Vertical Datum (NZVD) rather than stage heights. Yellow boxes show the NZVD elevation range of spawning sites discovered in April in four of these catchments. The length of the box represents the maximum window in which spawning could have occurred. Results from Oaro, Swan and Blue Duck clearly show the coincidence of spawning activity with periods of elevated water levels. At Harnetts this relationship is obscured by the presence of lower elevation spawning sites located downstream of the logger position combined with considerable gradient between the two. However, as with all other spawning sites observed in the study, all of the Harnetts Creek sites were also laid during periods of elevated flow. The prolonged periods of elevated water levels at Oaro and Kahutara result from interactions between river flows and barrier formation at the rivermouth. Similar effects can be seen at Swan in consideration of water level before and after the April rain events.

By late March, the continuing lack of rain suggested that the February eggs were unlikely to be re-inundated naturally and an ecological engineering experiment was designed in collaboration with the local council involving a temporary (4 hour) closure of the Lyell Creek mouth using a gravel bund sufficient to raise water levels upstream by approximately 40 cm. In addition to facilitating the hatching of February eggs, the March spawning surveys showed that this artificial inundation event triggered spawning in the affected reach despite its short duration. In contrast, no March spawning was recorded in any of the other study catchments (Fig. 2). However, a series of moderate rain events occurred in April sufficient to raise water levels region-wide, and spawning was detected in five of the study catchments. Concurrent water level and spawning site elevation data from the April events provide conclusive evidence of the triggering effect of water level fluctuations (Fig. 5b). Despite marked differences in the hydrographs, all spawning sites were located in an elevated position relative to the water level during the previous period of low flows.

These observations also provide insights into the likely duration of spawning events as revealed by the time at which water levels dropped to below the minimum elevation of spawning grounds. For example, the developmental stage of eggs at Blue Duck Creek suggested that spawning occurred on the 7 April rain event and the vertical position of eggs indicates that the spawning site was inundated for around 54 hours (Fig. 5b). In contrast, the mean spawning site elevations at Lyell Creek were 10 cm lower than the February sites but still a minimum of 20 cm above typical low flow levels. Together with the obviously fresh (non-eyed) appearance of the eggs, the elevation range indicates that spawning occurred during the relatively sharp spike in water levels on 22 April, for which the period of sufficient inundation lasted only a few hours. Harnetts Creek showed a similar pattern with spawning occurring close to the peak of the 22 April rain event within a short time period. These temporal patterns suggests that the fish are both highly attuned to detecting water level fluctuations and likely to be highly mobile within the river at these times. The surprisingly short lag times between the onset of water level fluctuations and spawning events indicate very rapid responses to environmental cues. Further research on these aspects is both intriguing and insightful for improving the understanding of adult fish movement and migration triggers. As shown here, these have a fundamental influence on spawning behaviour and the associated biogeography of spawning grounds.

## 4. Discussion

This study adds to previous evidence of plasticity in *G. maculatus* life histories (Augspurger et al. 2017), and is consistent with the prediction of Barbee et al. (2011) that the substantial differences in environmental characteristics found across the *G. maculatus* range could manifest as variability in spawning and migration cues. Moreover, recognition of diverse life histories in several amphidromous species has led to calls for reorientation of the term towards relationships between benthic and pelagic life stages rather than necessarily involving a marine-freshwater distinction (Closs et al. 2013; Closs & Warburton 2016). This highlights the need for further research on the characteristics of recruitment sources, and in turn emphasises the need for a solid understanding of the biogeography of spawning grounds which are the ultimate larval source (McDowall 2010b).

In New Zealand, there is a noticeable information gap on the characteristics of non-tidal rivers despite a considerable body of work on *G. maculatus* spawning in estuarine locations (Hickford & Schiel 2011; McDowall 1968, 1991; Orchard et al. 2018a), and contrasts with numerous studies on landlocked *G. maculatus* populations in Australian and South American rivers and lakes (Barriga et al. 2007; Chapman et al. 2006; Cussac et al. 2004; Pollard 1971). Recent advances in New Zealand have included the identification of non-diadromous recruitment in amphidromous galaxiid species, including *G. maculatus*, where suitable freshwater pelagic habitat was present in rivers open to the sea (David et al. 2019; Hicks et al. 2017). However, despite confirming the potential for spawning in non-tidal reaches of New Zealand’s river systems, these studies provide only limited information on the specific location and relative importance of the operative spawning sites. This is in part due to the choice of sampling methodologies which take advantage of otolith microchemistry markers as an indicator of previous life histories such as exposure to marine environments or other chemically-distinct water bodies such as lakes (Ruttenberg et al. 2005; Warner et al. 2005). Consequently, these approaches are well suited to identifying broad-scale phenomena such as the presence of non-diadromous individuals, but may lack the resolution needed to ascertain finer-grain aspects such the contribution of recruitment from different sources, and the associated biogeography of spawning grounds (Carson et al. 2013).

This study extends and complements the previous New Zealand work by applying a census survey approach with a focus on the characterisation of spawning grounds (Orchard & Hickford 2018). Although such surveys are resource-intensive in comparison to sparse sampling designs, they have the potential to elucidate fundamental spatial ecology aspects and generate information that is directly useful for site-based management needs. Findings from this study provide the first comprehensive account of spawning ecology in amphidromous *G. maculatus* populations at non-tidal rivermouths and include strong spatiotemporal patterns that were consistent across study catchments and months. The following sections provide a brief discussion of these aspects with a focus on transferable learning for conservation in other non-tidal rivers, and comparison with the existing knowledge base.

### 4.1 Position in catchment

The observed biogeographical pattern provides strong support for the prediction that spawning grounds would be found in non-estuarine lagoon environments due to similarities with known spawning areas in the lower reaches of tidal waterways. Five of the seven catchments featured a dynamic lagoon feature near the rivermouth, and in each case spawning was found within the lagoon or a short distance upstream. In the remaining two catchments (Blue Duck and Harnetts), no lagoon feature was present during the study but the spawning grounds were located in first slow-flowing reach upstream. At the catchment scale, all spawning grounds occupy a relatively compact reach within the lower river and in all cases high quality un-used spawning habitat was present further upstream. Although this pattern supports the notion of downstream fish movement to a preferred low-elevation and low-gradient reach as occurs in tidal situations (McDowall 1968), there was also some variation in the observed upstream limit of spawning between events. For example, in Lyell Creek in March, and Harnetts Creek in April, spawning was found further upstream of the previous records which translated to a larger confirmed spawning reach than if these sites had not been observed. Consequently, different river conditions and potentially also fish-centric factors such as the size of the spawning population could introduce further variation and result in a larger reach being important at other times. Despite this, the downstream limit of spawning was relatively consistent and strongly related to the most downstream position of suitable vegetation in the periodically inundated riparian zone (Fig. 3). This suggests that fish are actively searching the lower reaches of the river during spawning events and are selecting appropriate spawning habitat in that vicinity in preference to similar habitat further upstream.

In comparison to other studies, these results show similarities with previous reports of *G. maculatus* spawning in tidal lowland lakes such as Te Waihora / Lake Ellesmere in Canterbury, and lakes Waihola and Waipori in Otago, where spawning sites have been reported near the confluence of in-flowing streams (McDowall 1968; Richardson & Taylor 2002). In some cases, spawning has been recorded out-of-phase with the tidal influence in response to other sources of water level fluctuations that include wind-driven variations in water level height (Richardson & Taylor 2002). However, studies from lake environments in other countries have reported a variety of spawning behaviours including upstream migrations to spawning grounds in tributary streams (Pollard 1971), and lacustrine spawning in landlocked lakes (Barriga et al. 2002; Chapman et al. 2006). These variations demonstrate the need for further characterisation of spawning ecology across a range of environments and also the important role of adult fish migrations that are inherent in the selection of spawning sites.

### 4.2 Vertical dimension

Strong structuring in the vertical dimension was a common feature of all spawning sites (Fig. 4). At the reach scale, this resulted in the spawning grounds being 0.4 – 1.0 m above normal low flow levels and translated to a wide variety of positional nuances in relation to human activities in and adjacent to the river bed. At Blue Duck, Harnetts, Swan and Middle creeks the spawning grounds are located on relatively steep banks close the river channel where they are relatively protected from nearby recreational activities and stock grazing in adjacent land. However, at Lyell Creek which features well-defined banks in a more urban context, the spawning grounds overlap with areas subject to herbicide spraying and landscaping works. At Kahutara and Oaro rivers which feature wider river beds and more extensive rivermouth lagoons, spawning grounds are exposed to river diversion earthworks for flood control, vegetation clearance and off-road vehicle use.

These results have important consequences for river management since many human activities are found in the same elevation zone at these rivermouths, as is the case in tidal waterways (Orchard & Hickford 2018; Orchard et al. 2018a; Richardson & Taylor 2002). In the non-tidal context, the major differences include the lack of a regular (i.e., tidal) inundation cycle. This is generally useful from a management perspective by providing a convenient indicator of the most likely elevation range in which spawning will be found, leading to planning approaches based on tidal height as a proxy for the confirmed location of spawning sites (Greer et al. 2015). Despite this, multi-month surveys in tidal river systems have also shown that high discharge events can have the effect of raising spawning grounds above normal tidal levels, and this may be associated with appreciable horizontal translation away from the river channel into vulnerable areas such as recreation reserves (Orchard 2019). Consequently, attention to the role of discharge rates in the horizontal and vertical structuring of spawning grounds is important. This provides a sounds basis for the integration of flood management, vegetation control, and bankside recreational activities to achieve effective conservation at all rivermouths where *G. maculatus* populations are found.

### 4.3 Timing of spawning, fish movement and environmental cues

The timing of spawning in all seven catchments provided additional support for the hypothesis of spawning events being triggered by rain-induced water level fluctuations. Multiple lines of evidence including high precision elevation data and concurrent water level data for independent spawning events confirm the importance of this environmental cue with no exceptions across all 35 individual spawning sites. This functional relationship is consistent with *G. maculatus* ecology in tidal situations (McDowall 1968), and in landlocked riverine populations in which spawning was found in riparian vegetation inundated by freshes following heavy rain events (Pollard 1971). Insights on the timing of spawning events obtained from fine scale spatiotemporal data also suggest an important role for poorly understood aspects involving fish movement and migration triggers that precede the spawning event. The finding of a very short time lag between manifestation of the environmental cue and the onset of spawning deserves particular attention since this may be indicative of rapid fish movements at these times.

Results from this study show general support for the expectation that lunar cycles associated with spring high tide periods would be unlikely to act as a trigger for non-tidal spawning migrations, as has been reported to be a potential trigger in tidal waterways (McDowall 1968; Taylor 2002). However, it must be noted that the full moon period coincided with the heavy rain event of late February and could therefore have had a bearing on the position of the fish within the wider catchment at that time (Table 2). In comparison, some of the April spawning events show divergence from the lunar cycle, and an apparent synchrony with water level changes. Nonetheless it remains difficult to disentangle the major factors influencing fish movement towards rivermouths prior to the timing of spawning events, especially on a seasonal basis. For example, lunar cues might still be functioning as an influence on fish movements at the catchment scale in connection with progression of the *G. maculatus* lifecycle within these waterways.

Questions surrounding the functional trigger for fish movements in these non-tidal rivers add intrigue to the already mysterious process by which tidal river fish initiate their migrations to estuarine spawning grounds to coincide with spring high tides (McDowall 1968, 1991). Understanding these movements is of fundamental importance for sampling approaches that assume comparable study areas in relation to size-class distributions (Ravn et al. 2018), or in the case of mark-recapture techniques, require accounting for within-population detectability differences over time (Lindberg & Lindberg 2012; MacKenzie et al. 2003). For *G. maculatus*, the potential for variable dynamics between size or age classes combined with strong directional effects present confounding factors that are difficult to control for, yet could directly influence size-frequency and catch-per-unit-effort (CPUE) measures over relatively short time periods, including successive sampling days.

### 4.4 Protected areas implications and concluding remarks

Spawning grounds provide an example of a geographical small but disproportionately important critical habitat that is vital for successful completion of the whitebait life cycle (Mitchell 1994). Protecting spawning grounds is therefore an important focus for management, and is indeed a legislative requirement in New Zealand under the Conservation Act (1987) and Resource Management Act (1991). Spatially-explicit information on actual or likely spawning locations can assist these objectives by identifying areas for protection and enabling robust impact assessments to assess potential threats and determine management needs. For *G. maculatus*, spatial planning at relatively fine-scales has the potential to help address the common occurrence of riparian and river management activities that are incompatible with habitat protection goals. Applications include the avoidance of unnecessary trade-offs as a strategy to promote the efficient design of protected areas and related conservation tools (Faith & Walker 2002; Orchard & Hickford 2020). This study supports the quest for integrated management that embraces these aspects by providing the first comprehensive account of *G. maculatus* spawning ecology at non-tidal rivermouths and identifying biogeographical patterns that were consistent between rivers and over time. It is hoped that this may assist the development of both of regulatory measures and voluntary approaches to protect sensitive areas at the necessary times.

## 5. Acknowledgements

We thank the Ministry of Business, Innovation, and Employment and the New Zealand Ministry of Primary Industries for funding support. Particular thanks to staff at the Canterbury Regional Council for helpful discussions and access to water level records, and to staff and students at the University of Canterbury who supported this work.

## Supplementary material

**Fig. S1.**
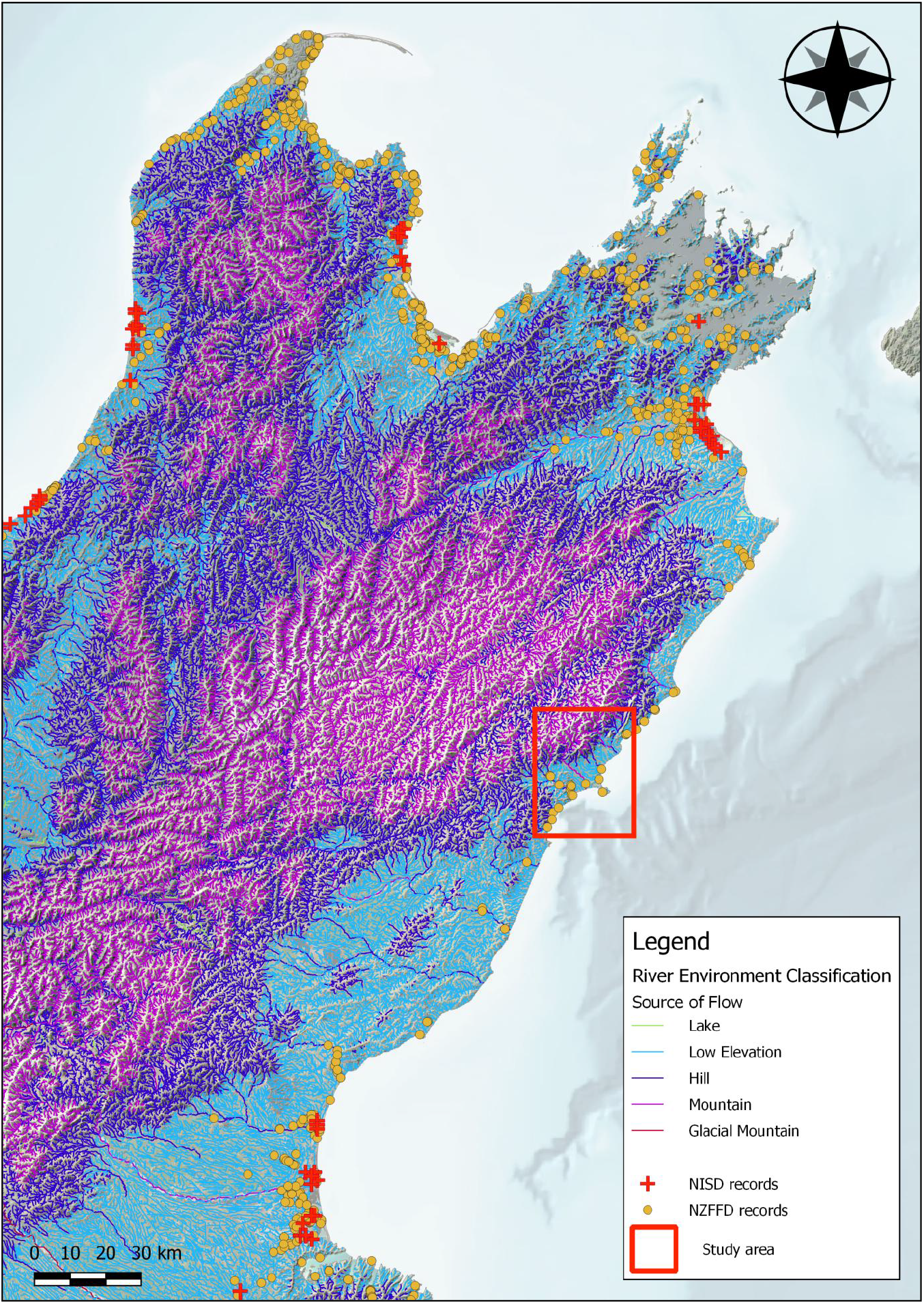
New Zealand Freshwater Fish Database (NZFFD) and National Īnanga Spawning Database (NISD) records for *Galaxias maculatus* in the northern half of New Zealand’s South Island, and study area on the Kaikōura coast. Note lack of spawning records in this region despite *G. maculatus* presence. Waterways are visualised according to source of flow data from the River Environment Classification (Snelder & Biggs, 2002).

**Fig. S2.**
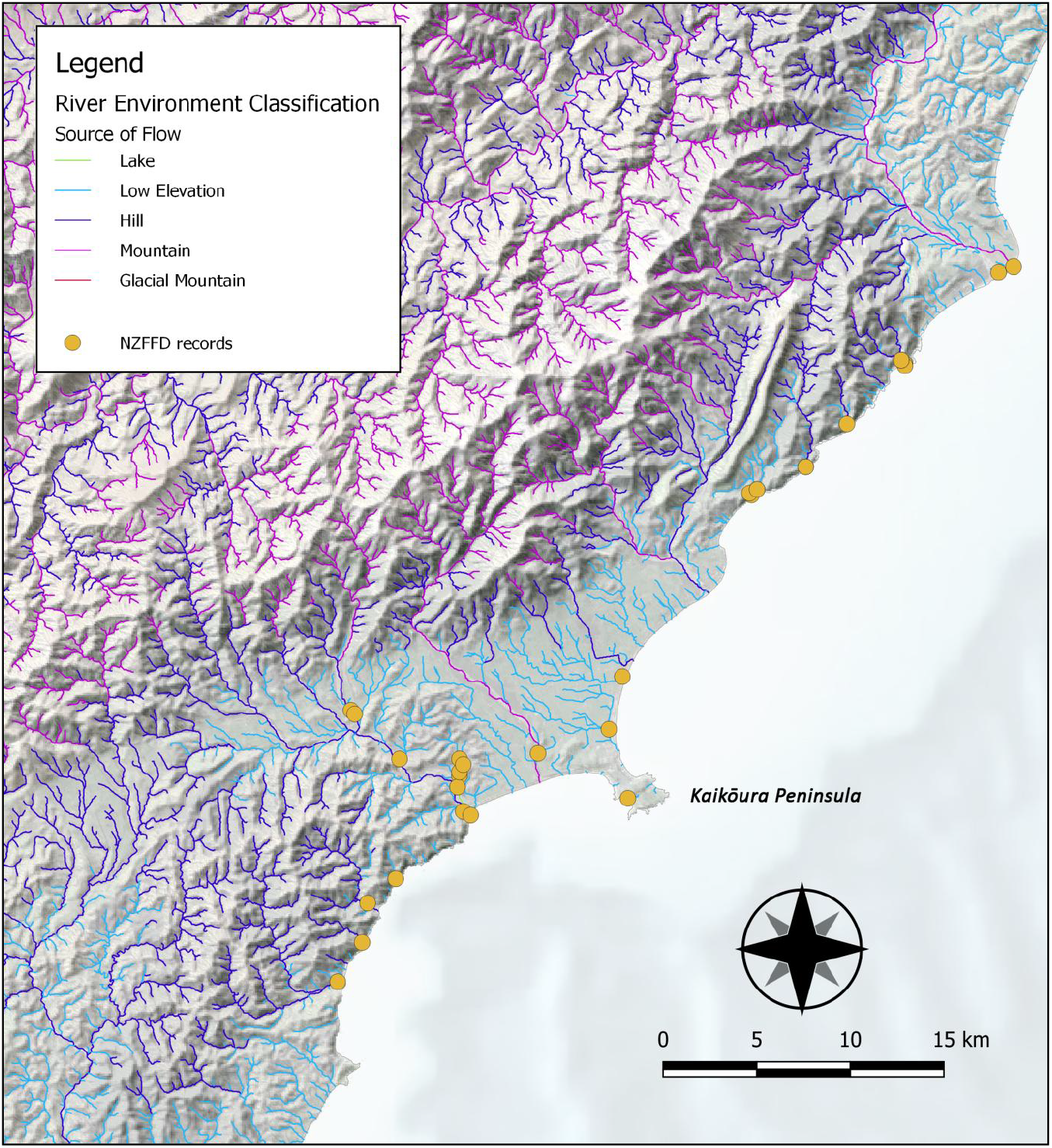
Closer view of waterways on the Kaikōura coast of New Zealand classified by source of flow (Snelder & Biggs, 2002). Point data are New Zealand Freshwater Fish Database (NZFFD) observations for *Galaxias maculatus*. Prior to the present study there were no National Īnanga Spawning Database (NISD) records or otherwise reported *G. maculatus* spawning sites within the map extent.

